# cran2crux: automatically create CRUX ports for R-packages

**DOI:** 10.64898/2026.05.09.723963

**Authors:** Petar B. Petrov, Valerio Izzi

## Abstract

**Motivation:** R together with CRAN and Bioconductor provides one of the richest ecosystems for bioinformatics and computational biology, with thousands of specialized packages. While GNU/Linux is a vastly-used operating system in this field, R-packages are typically managed independently of the system’s native package manager. This separation makes installation, updates and mass rebuilds cumbersome. CRUX, a minimalist semi-source GNU/Linux distribution, offers great flexibility with its ports-based system for the seamless integration of R-packages with its native package manager.

**Results:** The hereby presented cran2crux tool automatically generates CRUX ports for packages from both CRAN and Bioconductor. It performs recursive dependency resolution, handles naming conventions, extracts dependencies information, and supports inclusion of optional dependencies. The tool also provides convenient functions for checking updates and regenerating outdated ports. It can generate over 140 ports for complex packages such as Seurat in approximately 11 seconds, dramatically simplifying the maintenance of large R-dedicated repositories on CRUX.

**Availability:** cran2crux is available under the MIT license at https://github.com/izzilab/cran2crux. As of now, more than 650 R package ports, generated with the tool, are available in the CRUX ports database.

## Introduction

The R [1] programming language (https://www.R-project.org) and its ecosystem are heavily used in data science, with particularly strong adoption in bioinformatics and computational biology. R, together with CRAN (https://cran.r-project.org) and the bioinformatics-oriented Bioconductor project (https://bioconductor.org/), form one of the dominant platforms for the analysis of high-throughput biological data. This is made possible by thousands of R-packages, specifically tailored for genomics, transcriptomics, proteomics, single-cell analysis and more.

GNU/Linux is vastly used in bioinformatics, due to its reliability, powerful command line shell and the plethora of dedicated, open source tools. Software packaging is a core component of GNU/Linux distributions or any other UNIX-like operating system (OS). It provides a structured, centralized, and reliable way to install, update and remove software. R-packages are typically installed using R’s own package management tools (`install.packages()` for CRAN and `BiocManager::install()` for Bioconductor), placing them outside the system’s native package manager. Doing updates or mass rebuilds may, therefore, prove challenging and complicated, especially between major point releases of R,e.g. 4.5 to 4.6. Thanks to the flexibility of open-source, development of tools that bridge language-specific package ecosystems with the system-specific packaging is common.

CRUX (https://crux.nu/) is a minimalist and highly customizable GNU/Linux distribution, that offers an elegant ports system and an advanced ports managing tool, called `prt-get` [2]. The ports are simple text files (Pkgfile), using BASH syntax, facilitating users to create and maintain their own repositories (https://crux.nu/portdb/). Despite this transparency, creating and maintaining a large repository of interdependent ports -- such as one containing hundreds/thousands of R packages -- can quickly become tedious and time-consuming.

To simplify installing, updating, and managing R-packages on CRUX, we developed `cran2crux` -- a simple tool that automatically generates ports for CRAN and Bioconductor entries, with recursive dependency resolution. Although initially created to support bioinformatics-focused R-packages, `cran2crux` is designed to work with virtually any package from CRAN or Bioconductor.

### cran2crux

The core components of `cran2crux` are written in R and are invoked from a BASH script, that provides convenient command line options for ports generation and management. The code is freely available under the MIT license at Izzi Lab’s GitHub repository (https://github.com/izzilab/cran2crux). Although `cran2crux` supports both CRAN and Bioconductor, it is named in homage to `cpan2crux` -- a CRUX tool that performs a similar function for Perl modules.

When generating a port for an R-package named “Foo”, `cran2crux` adds the “r4-” suffix and converts the name to lowercase, resulting in “r4-foo”. Following CRUX packaging conventions, any dots in the name are replaced by dashes and any dashes in the version are replaced by dots, hence the produced package will be called, for example `r4-foo#1.0.0.tar.xz`. The package is then ready to be installed system-wide, using CRUX’s `pkgutils` [3], making it available to all users.

A configuration file is installed in `/etc` containing maintainer information, the preferred CRAN mirror, and the target Bioconductor version. When writing the port’s Pkgfile, `cran2crux` adds dependencies information from the *Depends, Imports*, and *LinkingTo* fields provided by upstream, while optional dependencies are retrieved from the *Suggests* field. Dependencies specified in the *SystemRequirements* field fall outside the R ecosystem and at the moment `cran2crux` is not meant to handle them. However, adding such functionality is considered for the future.

Generating a single port takes the following form:

~~~
$ cran2crux Foo
~~~

Parsing the `-r`(`--recursive`) option will generate ports for the R-package, it’s dependencies and their own dependencies, in a recursive manner:

~~~
$ cran2crux Foo -r
~~~

The tool first synchronizes with CRAN and Bioconductor, then recursively calculates dependencies (default depth of 5 iterations) before writing the ports to disk. The default iterations value rarely needs to be changed and may be left unspecified by the user, as in the example above. Alternatively, the `-ro` (`--recursive-opt`) argument can be used to also generate ports for optional dependencies (those listed in the *Suggests* field). Because this process is fully recursive too, it often results in the creation of thousands of ports and can be computationally intensive. For this reason, the `-ro` option should be used with caution. When used, the recursion depth can be increased by passing a *positive integer*, larger or equal to 2 (e.g. 10) after the `-r` or `-ro` option:

~~~
$ cran2crux Foo -ro 10
~~~

To view which installed packages have newer versions available upstream, the `-so` (`--show-old`) argument is passed to the script directly:

~~~
$ cran2crux -so
~~~

Finally, to generate ports updated to the newer versions, use the `-u` (`--update`) option:

~~~
$ cran2crux -u
~~~

### Example usage

As a demonstration, let’s create a port for KEGGgraph [4, 5], (an R-package providing a graph approach to KEGG), and its dependencies. KEGGgraph is available on Bioconductor (https://bioconductor.org/packages/KEGGgraph), while its dependencies are provided by both Bioconductor and CRAN. Generating ports will have the following output in terminal, listing the upstream repository, R-package name and the corresponding port that is created (e.g. r4-kegggraph):

~~~
$ cran2crux KEGGgraph -r
[ BioC ] Created port for BiocGenerics : r4-biocgenerics
[ CRAN ] Created port for bitops : r4-bitops
[ CRAN ] Created port for generics : r4-generics
[ BioC ] Created port for graph : r4-graph
[ BioC ] Created port for KEGGgraph : r4-kegggraph
[ CRAN ] Created port for RCurl : r4-rcurl
[ BioC ] Created port for Rgraphviz : r4-rgraphviz
[ CRAN ] Created port for XML : r4-xml
~~~

In this way, we have been maintaining an up-to date CRUX repository of over 650 ports, called “r4” (https://crux.nu/portdb/?a=repo&q=r4) for R-packages needed in our work, when doing software development in R. Among them are the Matrisome AnalyzeR [6] -- a suite to annotate and quantify extracellular matrix (ECM) molecules in big datasets across organisms (https://matrinet.shinyapps.io/MatrisomeAnalyzer), MatriCom [7], a tool which does single-cell RNA-sequencing data mining to infer cell–extracellular matrix interactions (https://matrinet.shinyapps.io/matricom), MatriSpace [8] -- an instrument for the identification and visualization of spatially resolved ECM gene expression patterns in health and disease (https://matrinet.shinyapps.io/matrispace), and ProToDeviseR [9] -- an automated protein topology scheme graphics generator (https://matrinet.shinyapps.io/ProToDeviseR).

Checking for potential updates of installed R-packages (such as the case of out >650), will report the port name, the R-package name, the currently installed version and the upstream version, as well as the repository itself:

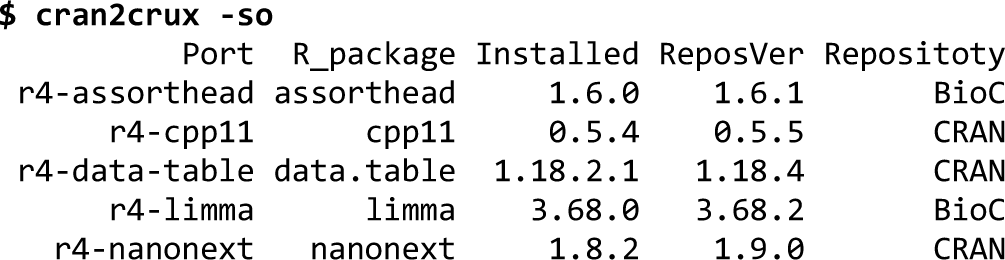

Then, generating updated ports is quite straightforward:

~~~
$ cran2crux -u
[ BioC ] Created port for assorthead : r4-assorthead
[ CRAN ] Created port for cpp11 : r4-cpp11
[ CRAN ] Created port for data.table : r4-data-table
[ BioC ] Created port for limma : r4-limma
[ CRAN ] Created port for nanonext : r4-nanonext
~~~

All that’s left to be done is to update the r4 CRUX repository, followed by a system update via `prt-get sysup` .

### Performance benchmarks

The Seurat package [10] is widely used in bioinformatics, and has been established as a standard tool for single cell transcriptomics analyses. It is available on CRAN (https://cran.r-project.org/web/packages/Seurat), where it lists over 50 first-level dependencies (*Depends, Imports* and *LinkingTo* fields). Running `cran2crux Seurat -r` generates total of 142 ports in about 11 seconds on an Intel i7-9700KF @3.6GHz, as reported by `time` . As a comparison, `cran2crux` was run by skipping the syncing step (data already downloaded beforehand) or setting the number of dependencies depth iterations to a lower value, e.g. 2 (Figure 1).

**Figure 1.**
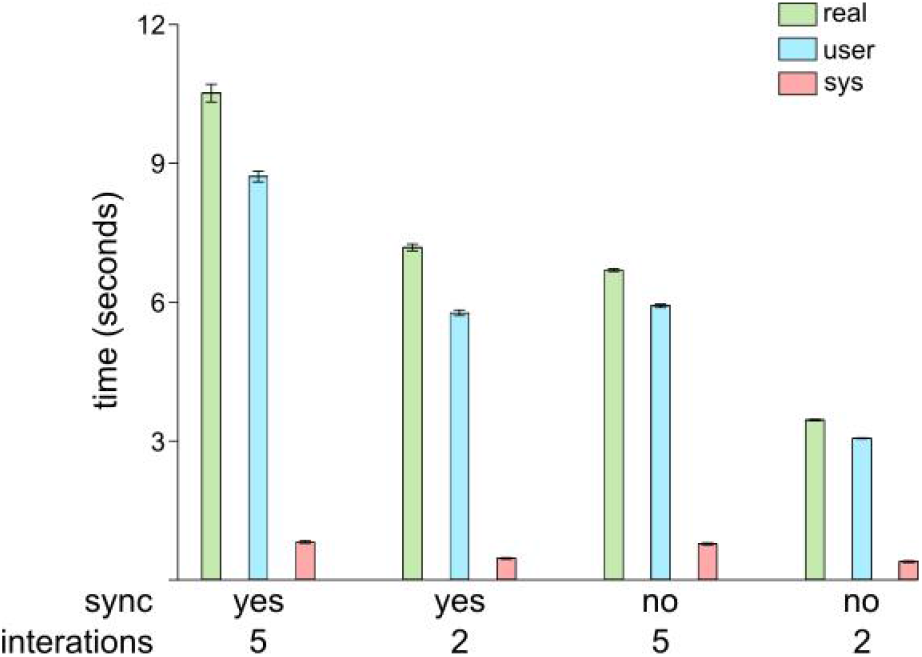
Performance benchmarks of cran2crux. Runs were timed with (default) or without CRAN and Bioconductor synchronization, as well as, with 5 (default) or 2 iterations for dependencies resolution. Error bars indicate standard deviation.

Three runs were timed for each combination of conditions and the mean values of *real* (total elapsed time), *user* (time in user-space code) and *sys* (time in kernel/system calls), were calculated and plotted in Gnumeric spreadsheet (https://gnome.pages.gitlab.gnome.org/gnumeric-web/). As expected, these run-times were shorter, illustrating that most time is spent on downloading package lists from upstream and resolving the full recursive dependency tree, while the actual local work on writing ports to disk is very fast.

### Adaptations to other distributions

Due to its open-source license (MIT) and relatively simple codebase, `cran2crux` can be easily ported or adapted to serve a similar purpose on other distributions. Any distribution that uses a ports-like packaging system — where each package is defined by a readable build recipe — could potentially benefit from a tool based on, or forked from `cran2crux` . Good candidates include Arch Linux (https://archlinux.org/), Slackware (http://www.slackware.com/) via the semi-official SlackBuilds.org project (https://slackbuilds.org/), Void Linux (https://voidlinux.org/) and Gentoo (https://www.gentoo.org/). A particularly interesting case is BioArchLinux [11], a community project that maintains a large collection of bioinformatics tools on top of Arch Linux using PKGBUILDs (https://bioarchlinux.org/). Since Arch Linux itself was originally inspired by CRUX, its PKGBUILD format is structurally very similar to CRUX’s Pkgfile. This makes BioArchLinux an excellent candidate for a `cran2crux` -based tool, as the core logic for generating package recipes would require only moderate adaptation.

## Conclusions

The hereby presented `cran2crux`, provides a fast and efficient way to generate and maintain large collections of R-packages in CRUX. Currently, there are over 650 R-packages ports available and actively maintained at the distribution’s ports database (https://crux.nu/portdb/?a=repo&q=r4). Thanks to its opens source license and simple code, it has the potential to be adapted to other UNIX-like systems that use a ports-like packaging.

## Acknowledgements

We thank the CRUX core team for their continued maintenance of the distribution and for keeping an always up-to-date port for R available.

## Author contributions

PBP wrote the code, performed the testing, did benchmarking and drafted the paper. VI acquired funding and contributed to the manuscript preparation.

